# The Foster method: Rapid and non-invasive detection of clinically significant American Foulbrood disease levels using eDNA sampling and a dual-target qPCR assay, with its potential for other hive pathogens

**DOI:** 10.1101/2021.06.18.449084

**Authors:** John F. Mackay, Rebecca E. Hewett, Noa T. Smith, Tammy L. Waters, John S. Scandrett

**Affiliations:** dnature diagnostics & research Ltd, 60 Carnarvon St., Gisborne, New Zealand; Scandrett Rural Ltd, Invercargill, New Zealand

**Keywords:** honeybee, American foulbrood, entrance, qPCR, diagnosis, swab, quantification, non-invasive

## Abstract

Clinical signs of American Foulbrood (AFB) can be difficult to diagnose and thus disease is often missed and leads to further spread. Diagnosis is centred on the beekeeper’s skill in recognising clinical symptoms – a highly subjective and time-consuming activity. Previous testing methods have relied on sampling that necessitates dismantling the hive and/or requires multiple visits to retrieve passive samples. The Foster method is a novel environmental DNA sampling method using colony entrance swabs together with a dual-target qPCR for *Paenibacillus larvae*: the causative bacteria of AFB disease. The quantification data generated can be used to detect hives with clinically significant infections, even before visual symptoms are apparent. Such a sampling method will be applicable to other bee pathogens and incursion pests.

**Importance:** Discovery of American foulbrood disease in a honeybee colony typically means the destruction of the bees and hive by burning, in New Zealand and many other countries. This discovery is typically by visual examination of capped brood by the beekeeper - a subjective skill that means the disease is being missed or not recognised. It is a time-consuming and exacting process to inspect hives for AFB. Here we present a novel rapid sampling method that does not require opening/ dismantling the hive, in conjunction with a dual-target quantitative PCR assay for the bacteria responsible, *Paenibacillus larvae*. Using the resulting quantitative data, hives presenting visual clinical symptoms or likely to soon become clinical, can be determined and the hives dealt with appropriately before further spread occurs. This study provides the basis for a novel way of sampling for honeybee pathogens and pests.

## Introduction

American Foulbrood (AFB) is one of the most destructive diseases in honeybees (*Apis mellifera*) and is caused by spores of the Gram-positive bacteria *Paenibacillus larvae* infecting bees during their larval stage. These spores are extremely hardy; being resistant to heat, caustic, and other chemical treatments with the ability to remain infectious for over 30 years (Genersch, 2010). Only a few spores are required to cause an infection in a new larva and yet a single infected larvae can generate many millions of spores - leading to rapid spread and decline in a bee colony. Once the colony has weakened or died then it may be ‘robbed’ by bees from neighbouring hives where contaminated honey is brought back to a neighbouring hive, which leads to further infections. Infection is also spread unknowingly by the beekeeper (Fries and Camazine, 2001), through the introduction of hive frames carrying an undetected infection into a new colony of bees or the feeding of contaminated honey to colonies.

In New Zealand, hive numbers have more than doubled since 2013 (www.afb.org.nz). Similarly, the number of beekeepers has also increased – primarily due to hobby and conservation efforts (several hives per beekeeper) but also in the commercial sector (hundreds or thousands of hives per operation). Much of the growth in hive and commercial beekeeping has been due to the increased demand for high-value New Zealand manuka honey. The increase in hive numbers (and densities) combined with newer beekeepers, who are less likely to be skilled in identifying the clinical signs of AFB, has led to an increase in the percentage of hives reported by beekeepers as infected from 0.21% in 2015 to 0.46% in 2021 (www.afb.org.nz).

The diagnosis of AFB is based primarily on visual symptoms – typically unusual, capped brood cells on a beehive frame that warrant investigation through a ‘roping’ field test. In a hive with three to seven frames of brood or more, this means typically thousands of brood cells to visually examine per hive. Other diseases mediated by varroa can affect brood cells meaning that examining suspicious cells for clinical AFB signs is laborious and prone to disease being missed. Clinical signs can vary based on the AFB genotype present, with ERIC II strains exhibiting reduced clinical symptoms (Genersch, 2010). In addition, hives must be dismantled, and the colony disrupted to inspect for AFB.

Confirmation of AFB in beehive materials has been traditionally performed by culture, using selective media, and induced germination of spores. However, this is a variable process (Forsgren, 2008) and may take up to a week for results. PCR and qPCR (quantitative PCR) have been demonstrated as effective tools for the detection of *P. larvae*, with qPCR being the preferred diagnostic tool in recent years due its faster time to results and lower chance for amplicon contamination. However, most applications have not used the quantitative data generated; rather using the tool for rapid confirmation of culture isolates (Kňazovická et al, 2011). The quantification aspect is important as low levels of AFB are often not clinically relevant (Bassi et al., 2018, Pernal and Melathopoulos, 2006) and such hives may never develop clinical symptoms. Much of this low level may be due to the hygienic behaviour of bees to reduce overt infections (Spivak and Reuter, 2001).

Most molecular publications relating to AFB to date have relied on either conventional PCR or SYBR Green-based qPCR and the sole use of the 16S gene due to its high sensitivity as a multicopy target (Dobbelaere *et al.*, 2001; Piccini *et al.*,2002). However this sensitivity can be at the expense of specificity, as evidenced by the initial design (Martinez, 2010) and subsequent re-evaluation/re-design of qPCR primers to improve specificity (Rossi *et al.*, 2018). Sensitivity for bacterial targets can also be compromised by deletions of target sequences or sequence polymorphisms which affect primer or probe binding (Dahlberg *et al.*, 2018; Johansen *et al.*, 2019; Xiu *et al.*, 2021) and many microbiological qPCR tests now routinely utilise two independent targets to enhance both sensitivity and specificity, such as many of the commercial qPCR tests for chlamydia and the virus behind COVID-19: SARS-CoV-2. Hydrolysis probe (“TaqMan^®^”) qPCR assays permit quantification of each target by selected fluorescent wavelengths (Kušar *et al.*, 2021) while also allowing the use of an internal control in an additional detection channel for confirmation of suitable quality DNA present in the reaction. These internal controls may improve the sensitivity of previous studies whereby the presence of inhibitors possibly prevented the amplification of *P. larvae* targets. (Forsgren and Laugen, 2013)

We used a commercially available multiplex qPCR assay for *P. larvae* (Hall et al., 2021) and developed a new and rapid environmental DNA (eDNA) sampling method for the detection of hives having - or at risk of developing - clinical AFB infections. Previously *P. larvae* has been detected in debris from a hive baseboard (Bassi et al., 2018) but this study required installing collection sheets, necessitating dismantling the hive twice as well as repeat visits. The ability to detect relevant levels of *P. larvae* from a single sample taken from the entrance of a hive would permit far more rapid sampling of hives and allow infected hives to be dealt with more quickly (Lyall et al., 2019). In this study, we evaluated and compared the levels of *P. larvae* among bees, the entire baseboard, and the entrance region of the baseboard.

## Methods

Quarantining hives in an apiary upon discovery of a clinical AFB hive is a common international practice, in order to minimise spread of the disease. From five such quarantine apiaries around New Zealand (East Coast, Wellington, Wairarapa and South Canterbury regions) multiple hives were sampled two-monthly from late 2018 to early 2020. In addition, samples from visually clinical hives and neighbouring hives were collected by local apiary inspectors for testing during this time. Samples included nurse bees, swabs from the hive entrance and swabs from over the entire baseboard. Subsequent samples comprised of comparisons between bees and solely the hive entrance. Bees were collected from the brood frames using 50 mL sterile containers and frozen to euthanise the bees. Sterile foam-tipped swabs (Puritan) were moistened in sterile water before swabbing either the whole baseboard or just through and across the hive entrance for three to five seconds. Instructions were provided to participating beekeepers to rotate the swabs during collection to ensure all swab surfaces could collect hive material. Swab heads were then snapped off into a 2 mL microcentrifuge tube, capped and sent to the laboratory with the bee samples, at ambient temperature by overnight courier. Once in the laboratory, bee samples were frozen at −20°C and swabs were stored at ambient temperature until DNA extraction was performed.

Genomic DNA extraction from bees and associated bacteria was performed from each sample containing 10 bees. DNA was extracted using the Bee Pathogen DNA/RNA Extraction Kit (dnature diagnostics & research Ltd, Gisborne, New Zealand); a column-based kit in conjunction with beadbeating. 10 bees were used for the practical purpose of ease of sample handling and throughput. In short, 3 mL DXL lysis buffer supplied in the kit was added to 10 bees in a 5 mL tube, containing a mixture of 0.5 mm (~2.4 g) and 2.3 mm zirconia silica (~1 g) beadbeating beads (BioSpec Products, Bartlesville, USA). The bees were homogenised in a Mini Beadbeater 16 instrument (BioSpec Products, Bartlesville, USA) for 3 minutes. Tubes were then incubated at 65°C for 10 minutes with manual inversion every 3 minutes. 1 mL of the homogenate was transferred to a 2 mL microcentrifuge tube and centrifuged at 15,000 x g for 5 minutes. 500 μL of supernatant was transferred to a new microcentrifuge tube containing 450 μL AD buffer and mixed well. The solution was then applied to a nucleic acid binding column from the kit (in two passes) and centrifuged through before successive washing steps and elution of nucleic acids from the column in 100 μL elution buffer.

A similar process was used for swabs: they were placed in a screwcap 2 mL tubes with 0.5mm zirconia silica beads (~0.8g) and 2.3 mm zirconia silica beads (0.3g). 800 μL DXL buffer was added to each tube and the swab material homogenised in the Mini Beadbeater 16 for 3 minutes. The tubes were then incubated at 65°C with shaking for 10 minutes, before centrifuging and processing 500 μL of the resulting supernatant as for the bees. For pooled applications, individual swabs were added to 500 μL DXL buffer in a separate 2 mL microcentrifuge tube and vortexed for 10 seconds to dislodge cells and bacteria off the swab. 60 μL of each swab eluate (up to twelve swabs) was added to a 2 mL beadbeating tube as for single swabs and processed as for single swabs from the beadbeating step.

Assessment of DNA quality and presence of potential inhibitors was performed by measurement (spectroscopy) on a DS-11 FX (DeNovix, Wilmington, USA) and OD260/280 measurements recorded. Dilutions of extracted DNA were also tested by qPCR to measure the expected Cq increases with increasing dilution and thus monitor any qPCR inhibition.

The AFBduo qPCR test (dnature diagnostics & research Ltd, Gisborne, NZ) is a commercially available multiplex hydrolysis probe assay that uses two targets for *P. larvae:* a single copy gene for quantification purposes (ftsZ, a prokaryotic homologue to tubulin) and the multicopy 16S gene. The ftsZ hydrolysis probe is detected in the FAM channel and the 16S target in the HEX channel. The test also includes a simultaneous internal control hydrolysis probe reaction (ROX channel) for an animal mitochondrial marker, to confirm validity of results (in the cases where *P. larvae* was not detected) by way of successful internal control amplification.

Each AFBduo qPCR reaction was performed in a 10 μL volume and consisted of 5.5 μL PCR grade water, 0.5 μL of the 20X Oligo Mix and 2 μL of the 5X Mastermix (all reagents supplied in the AFBduo real-time PCR kit, dnature). 2 μL genomic DNA, positive control DNA (DNA extracted from larvae with clinical disease) or sterile water (for no template controls) was added per reaction. The amplifications were performed on the Mic qPCR Cycler (BioMolecular Systems, Australia) or the Eco qPCR instrument (Illumina, San Diego, USA) and the cycling conditions comprised: denaturation at 95°C for 2 minutes followed by 40 cycles of 95°C for 5 seconds and 60°C for 15 seconds for both instruments. Data was acquired on the Green (ftsZ), Yellow (16S) and Orange (Internal control) channels of the Mic cycler or FAM, HEX and ROX channels on the Eco instrument. Cq’s (Cycle quantities) were automatically generated by the Mic software, using the dynamic analysis setting (based on second derivative maximum method).

A synthetic DNA standard (gBlock, IDT, Singapore) was utilised that incorporated both amplicon sequences (available on request). The synthesised 153 bp standard was resuspended in TE buffer, pH8, quantified in nanograms per μL on a DS-11 FX microvolume spectrophotometer (DeNovix, Wilmington, USA) and its copy number calculated using the equation:

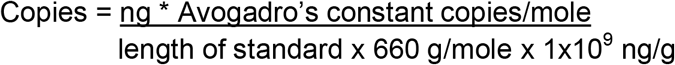

Quantification standards were generated from 10^6 copies per μL down to 2 copies per μL (Svec et al., 2015). For all dilutions, the diluent was a pooled DNA sample extracted from bees free of *P. larvae*. Dilutions of extracted DNA from samples positive for *P. larvae* were diluted in a pool of negative bee DNA.

The standards and their resulting Cq’s (supplementary S1) were also used to convert the Cq’s resulting from bee and swab samples to spore levels using the amplification efficiency of the single copy ftsZ gene, the y-intercept, and the formula:

Spores per swab or 10 bees = 10^(Cq ftzZ-39/−3.48)^ × 50

Statistical analysis was performed after imputing data points by regression analysis of the other paired data, for two clinical hives (due to spore numbers being skewed lower by the new hiveware baseboards at time of sampling) resulting in a total of 26 data points with paired clinical data. A binary logistic regression model was performed in Minitab (Minitab^®^, State College, PA, USA) to identify the relationship between spore numbers at the entrance and clinical disease likelihood, and analysis of variance calculated to confirm the correlation between increased entrance spores and the clinical signs of AFB. A *p*-value of <0.05 was considered significant.

## Results

Testing of the AFBduo assay on synthetic DNA standards showed high analytical sensitivity, with both target reactions in the assay able to reliably detect down to 4 copies per 10 μL reaction. The reaction targets amplified with high efficiency in the multiplex (ftsZ = 94% and efficiency for 16S = 96%) as well as excellent linearity over the 7 orders of magnitude (r = 0.998). On testing *P. larvae*-positive bee, honey, and hive material, the 16S target typically amplified 3-5 cycles earlier (i.e. lower Cq’s) than the ftsZ target as expected, due to the multiple copies of the 16S sequence per AFB bacterium compared to the single copy ftsZ gene.

The DNA extraction method from bees generated high purity DNA as shown by measurements on the microvolume spectrophotometer with OD260/280 measurements of between 1.8 and 1.9. Extracted DNA samples serially diluted in elution buffer showed the expected increase in Cq’s with decreasing concentration, indicating no inhibitors were present in the samples following DNA extraction. DNA yields from swabs were too low to be measured on the spectrophotometer, however internal control Cq’s for serial dilutions as for the bees, indicated the DNA was similarly of suitable quality.

In this study, we evaluated and compared the levels of *P. larvae* among bees, the entire baseboard, and the entrance region of the baseboard. In the first cohort of quarantine beehives from multiple regions in NZ (beehives from quarantine apiaries i.e. an apiary that had contained a hive exhibiting clinical signs), 45 hives were sampled up to 6 times during the 2018-2019/20 seasons. The samples taken were nurse bees, swabs from the entire baseboard and entrance swabs. Among these hives, 8 were found to have clinical signs of AFB (‘roping’ larval remains) during sampling while another 6 were presumed to have AFB due to their condition and the high prevalence in the apiary. Examples of the progression of *P. larvae* levels in three of the colonies is shown in Table 1, where the resulting Cq data has been converted to spores per 10 bee or swab sample processed.

**Table 1:** AFB spore levels seen in bee samples and entrance swabs from three quarantine hives examples followed over time. Total number of hives followed over time was thirty.

| Date | 14 Sept<br>2018 | 4 Dec<br>2018 | 16 Jan<br>2019 | Clinical status |
| --- | --- | --- | --- | --- |
| Bees<br>(spores/10 bees) | - | 24,448 | 779,266 | Clinical signs seen 16 Jan after<br>sampling and hive destroyed |
| Entrance<br>(spores/swab) | ND | 10,870 | 678,363 |  |

Hive STF4
| Date | 22 Sept<br>2018 | Clinical status |
| --- | --- | --- |
| Bees<br>(spores/10 bees) | 2,591,937 | Clinical signs seen<br>early October 2018<br>and hive destroyed |
| Entrance<br>(spores/swab) | 4,281,312 |  |

Hive RL02
| Date | 31 Oct<br>2018 | 27 Dec<br>2018 | 11 Mar<br>2019 | 23 Aug<br>2019 | Clinical status |
| --- | --- | --- | --- | --- | --- |
| Bees<br>(spores/10 bees) | ND | ND | ND | ND | No AFB seen |
| Entrance<br>(spores/swab) | ND | ND | ND | ND |  |

ND Not Detected
- Test not performed

In a second cohort of hives from one beekeeper, 23 hives were tested in a quarantine apiary using bee and hive entrance swab samples. Four of these hives showed clinical signs of AFB. Levels of AFB in nurse bee samples were compared to the levels indicated by the entrance swabs. Once again, most hives showed no or low levels of *P. larvae*, while those with clinical AFB demonstrated much higher *P. larvae* spore levels in both the bee and entrance swab samples as shown in Table 2.

**Table 2:** Testing an apiary comparing bee and entrance swab samples. The spore levels marked with * indicate low level hive entrance spore on colonies recently placed into new hiveware (hives 11 and 12 were started from splits of hive 10). While at the time of sampling hive 11 showed no AFB symptoms (**), it developed clinical symptoms 2 weeks later and was destroyed. ND Not detected

|  | Clinical signs of AFB? | Bees | Entrance Swab |
| --- | --- | --- | --- |
|  |  | Spores/10 bees | Spores/swab |
| **MLHive 1** | **Yes** | **37,097,589** | **3,138,994** |
| **MLHive 2** | **Yes** | **514,061** | **24,008** |
| MLHive 3 | No | ND | ND |
| MLHive 4 | No | 387 | 1,107 |
| MLHive 5 | No | 539 | 377 |
| MLHive 6 | No | ND | ND |
| MLHive 7 | No | 80 | ND |
| MLHive 8 | No | 210 | ND |
| MLHive 9 | No | 205 | ND |
| **MLHive 10** | **Yes** | **55,866,177** | **962,621** |
| MLHive 11 | No** | 9,518,751 | ND* |
| **MLHive 12** | **Yes** | **12,561,091** | **890\*** |
| MLHive 13 | No | <10 | <10 |
| MLHive 14 | No | ND | <10 |
| MLHive 15 | No | 2049 | 1,318 |
| MLHive 16 | No | ND | 861 |
| MLHive 17 | No | ND | 326 |
| MLHive 18 | No | ND | 1,092 |
| MLHive 19 | No | 97 | 207 |
| MLHive 20 | No | ND | 11,235 |
| MLHive 21 | No | ND | 326 |
| MLHive 22 | No | ND | 201 |
| MLHive 23 | No | 943,739 | 2,365 |

A third cohort of hives came from a beekeeper with a high incidence of AFB that was introduced through the purchase of contaminated equipment into his operation. Here we performed initial trials on pooling swab eluates into single extractions to reduce time and costs. Swab eluates were pooled into groups of eight and later twelve, the DNA extracted from the pooled sample and tested for the presence of P. larvae DNA. The remnant samples from pools that tested positive for both markers, were then tested individually as described. The results (Table 3) show that that even in the pools of twelve hive samples, single hives with higher levels of AFB could be clearly detected within a pool, prompting testing of those hives contributing to the positive pool to identify potentially clinically affected hive(s).

**Table 3:** composite pooling of swabs to rapidly find clinically relevant levels in hives. In this instance the first pool showed raised levels and the remnant swab eluates were tested individually, whereupon elevated levels were found in hive SB219. Subsequent hive inspections showed this hive to have clinical symptoms and it was destroyed. – test not performed

|  | Pooled result (12)<br>ftsZ spore levels | Individual Swab Result<br>ftsZ spore levels (per<br>swab) | Clinical Signs of AFB<br>(following testing) |
| --- | --- | --- | --- |
| Hive SB216 | **15,630** | Not detected | No |
| Hive SB217 |  | Not detected | No |
| Hive SB218 |  | Not detected | No |
| Hive SB219 |  | **165,103** | **Yes** |
| Hive SB220 |  | Not detected | No |
| Hive SB221 |  | Not detected | No |
| Hive SB222 |  | Not detected | No |
| Hive SB250 |  | Not detected | No |
| Hive SB251 |  | Not detected | No |
| Hive SB252 |  | Not detected | No |
| Hive SB253 |  | Not detected | No |
| Hive SB254 |  | Not detected | No |
| Hive SB255 | Not Detected | - | No |
| Hive SB256 |  | - | No |
| Hive SB257 |  | - | No |
| Hive SB258 |  | - | No |
| Hive SB259 |  | - | No |
| Hive SB260 |  | - | No |
| Hive SB261 |  | - | No |
| Hive SB262 |  | - | No |
| Hive SB263 |  | - | No |
| Hive SB264 |  | - | No |
| Hive SB265 |  | - | No |

Statistical analyses using a binary logistic regression model (Minitab) and analysis of variance (Supplementary figure 4) indicated the strong relationship between the increasing spore levels and clinical signs (p = 0.00). Indeed, in this work a spore quantity of 565,000 or more, revealed a 95% probability of clinical signs being present in the hive.

## Discussion

The AFBduo assay provided high confidence in the results due to the integrated internal control and detection of two markers for AFB. As expected, the 16S marker had higher analytical sensitivity (amplifying 3-5 cycles earlier) due to the multi-copy nature of this marker (Dingman, 2017). However, the variability in Cq difference between the two markers among samples indicated the 16S copy number possibly varied among AFB isolates and thus could not be relied on for quantification (Dahllof et al., 2000). This variability in copy number for diagnostic targets has been noted in other diagnostic qPCR assays for AFB (Rossi et al., 2018; Dainat et al., 2018). Therefore the use of the single copy ftsZ gene was used for quantification purposes using a standard curve prepared with a quantified synthetic template diluted in AFB-negative bee DNA.

Three sample types were compared in the first cohort – bees from the brood nest (nurse bees), swabs from the whole baseboard surface and swabs through the entrance of the hive only. The ability to sample the hive through the entrance removed the need to dismantle and disrupt the hive as with the first two sample types (bees and whole baseboard swabbing). The levels of both *P. larvae* and internal control DNA seen on the swabs taken from the whole baseboard or just the entrance area of the baseboard were similar (Table 4) possibly due to the maximum amount able to be collected on a given swab. This showed dismantling the hive to swab the whole baseboard was not required, in order to detect the higher levels of *P. larvae* in clinically infected colonies. While the spore levels varied among clinically affected hives, they were still easily differentiated from the very low or undetectable spore levels in non-infected hives. While spore levels never increased during the monitoring time for many of the hives or became undetectable in some low spore cases, some of the hives showed increasing levels at each testing point before clinical signs were observed in the respective hive and the hive destroyed (Table 1).

**Table 4:** Comparison of baseboard and entrance swabs showing the differentiation of spore levels among two clinical hives and a hive without clinical symptoms.

| Hive | Location | Estimated Spore level<br>(ftsZ target) | Hive status |
| --- | --- | --- | --- |
| ML Shearer | Baseboard | 52 million | Clinical signs of AFB |
|  | Entrance | 25 million |  |
| ML Ellis | Baseboard | 3.6 million | Clinical signs of AFB |
|  | Entrance | 3.4 million |  |
| FL 10 | Baseboard | 61 | No AFB seen |
|  | Entrance | 335 |  |

Spore levels were the highest in the bees from the brood nest area of the hive and while other studies have employed up to 100 bees per sample for culturing, the use of 10 bees permitted easier and more convenient handling for DNA extraction. Despite the lower spore amounts on swabs, the levels from the entrance swabs could clearly be differentiated from levels seen in hives not exhibiting any visual signs of AFB (Table 2). While most hives without clinical signs of AFB had low or undetectable levels of *P. larvae* DNA detected, one hive without visual signs of AFB (Hive 11) showed levels similar to two other hives with clinical AFB signs. Indeed, at the next inspection by the beekeeper (approximately two weeks after sampling), hive 11 showed clinical signs of AFB and was destroyed thus demonstrating the ability of qPCR to find infected hives that may have pre-clinical infections or have symptoms missed by visual inspection.

However these results also demonstrated the importance of knowing the history of the hives sampled. In two cases (hives 11 and 12 in Table 2), the bees showed high levels of *P. larvae* spores while the hiveware showed low or undetectable levels. Subsequent investigation showed that the hiveware was new (< 2 weeks old at time of sampling) for both these hives and that the colonies were newly established from an unknowingly infected parent colony. Therefore, while the bees carried high levels of *P. larvae* there had not been enough time for spores to be deposited around the hive entrance surfaces. Further research will be required to assess this accumulation rate of spores on the hive as well as further refine the clinical spore threshold levels as more hives are tested over time.

Given the ability to detect clinically relevant levels of AFB disease through this simple hive entrance swabbing method, preliminary work with these entrance swab samples indicate that the same DNA sample can be used to test for other bee pathogens. *Nosema ceranae* has been detected through the use hive debris and qPCR (Copley *et al.*, 2012) and showed good correlation of levels between bees and debris. Since the development of this current work, a report has used hive debris collected over the period of a week to estimate loading of European foulbrood (a disease not present in New Zealand currently). As with the results described here, the bees offered a higher sensitivity of pathogen detection, yet the hive debris results still provided predictive information as compared to non-infected hives (Biová *et al.*, 2021). Another application could for biosecurity surveillance for unwanted hive pests such as tracheal mites (*Acarapis woodi*), tropilaelap mites or small hive beetle (*Aethina tumida*). The use of the sampling technique here combined with an appropriate qPCR assay (e.g. Li et al., 2018; Ward *et al.*, 2007) could provide rapid detection of any incursion to countries such as New Zealand (where tracheal and tropilaelap mites as well as small hive beetle are not found) by providing faster sampling of hives than dismantling hives to extract hive debris for testing (Ward *et al.*, 2007). The quantification provides an important risk element of the hive as demonstrated - as to whether visual signs are likely present at the time of sampling or likely to demonstrate these clinical signs in the near future. *P. larvae* has been detected from winter debris previously using conventional PCR (Ryba et al., 2009) and thus the quantification described in this current work, was not possible. All the methods published to date have required either dismantling the hive to sample from the baseboard or repeated visits to the hive to insert and remove sampling sheets. In contrast the rapid sampling method described here - that we have named the Foster method - is a fast, one-time sampling and quantification method amenable to composite testing and one that provides clinically significant information as to the AFB status of the hive.

## Acknowledgements and memoriam

This work was partly funded by Eastland Community Trust (now Trust Tairawhiti) (JFM, REH., NTS.,TLW) and the Sustainable Food and Fibre Futures from Ministry of Primary Industries, Alpine Honey and NZ Honey Trust (JSS). We thank all beekeepers who provided samples. We also thank Richard Hall (Ministry for Primary Industries) for early discussions, Dr Graeme Wood for statistical assistance and also Prof. Phil Lester and Dr Bart Janssen for comments on the manuscript. The method is named for New Zealand beekeeper Barry Foster - a long-time AFB educator, inspector, and former New Zealand AFB management agency board member. This paper is dedicated to the memories of Russell Poulter, a long-standing collaborator who passed away suddenly in January 2021 and beekeeper/AFB inspector John Maynard who passed away suddenly in February 2021.

## Supplementary information

**S1.** Sensitivity of AFBduo qPCR assay Synthetic standard was diluted in a *P. larvae-free* hive swab DNA matrix from 10^6 copies down to 2 copies per μL. Dilutions were amplified in duplicate and the mean Cq values presented.

| **Standard**<br>**(copies/μL)** | **Cq**<br>**ftsZ (FAM)** | **Cq**<br>**16S (HEX)** | **Cq**<br>**IC (ROX)** |
| --- | --- | --- | --- |
| 1,000,000 | 17.06 | 15.6 | 31.14 |
| 100,000 | 20.73 | 19.20 | 30.15 |
| 10,000 | 24.33 | 22.77 | 29.42 |
| 1,000 | 27.84 | 26.22 | 29.15 |
| 100 | 31.29 | 29.55 | 29.15 |
| 10 | 34.94 | 33.17 | 29.36 |
| 5 | 36.06 | 34.07 | 29.03 |
| 2 | 37.08 | 36.38 | 29.39 |

**S2:**
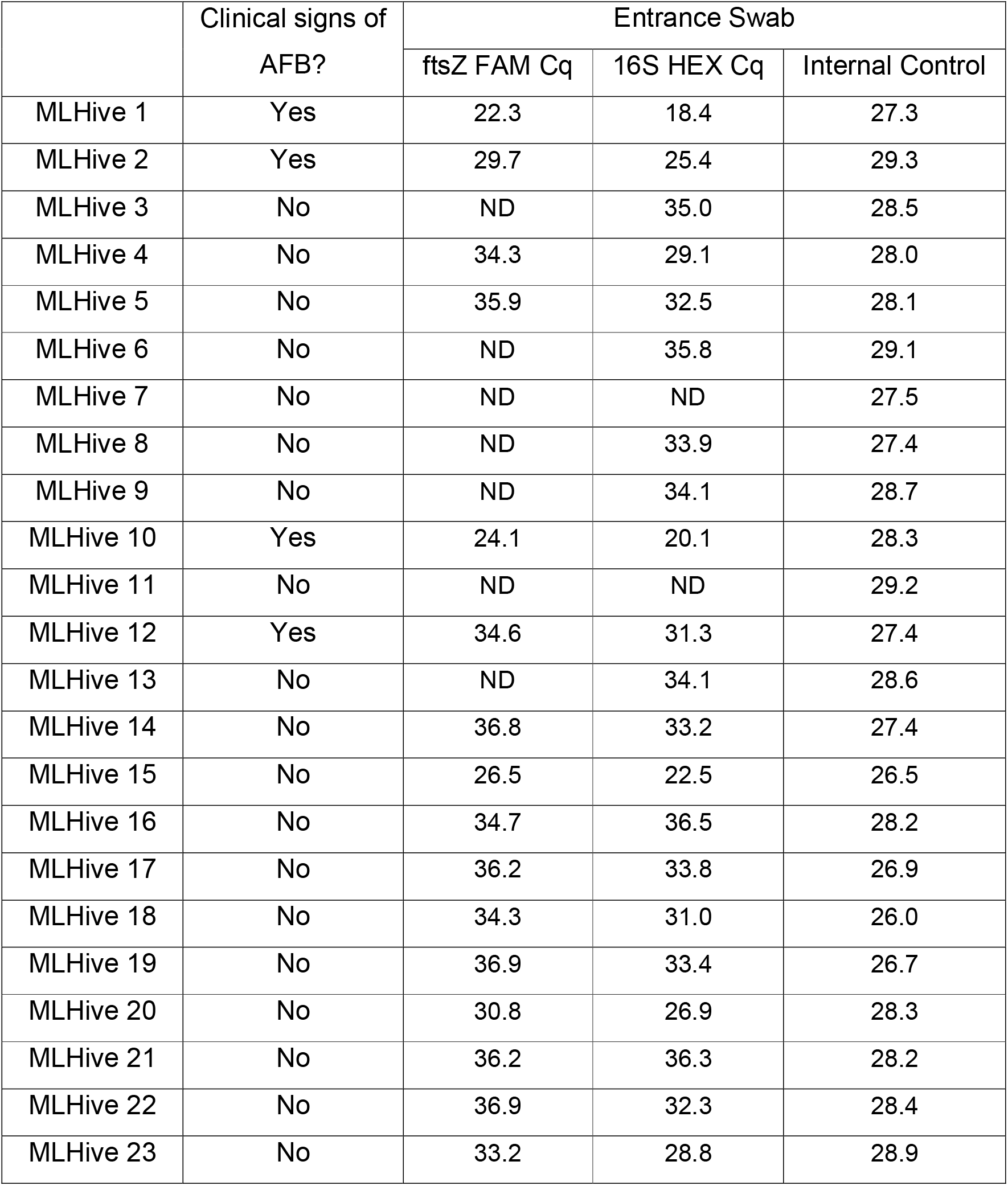
Cq data for hive series ML – *entrance swab* (spore levels in table 2)

**S3:** Cq data for hive series ML – *bees* (spore levels in table 2)

|  | Clinical signs of<br>AFB? | Bees |  |  |
| --- | --- | --- | --- | --- |
|  |  | ftsZ FAM Cq | 16S HEX Cq | Internal Control |
| MLHive 1 | Yes | 18.5 | 15.1 | 25.2 |
| MLHive 2 | Yes | 25.0 | 21.0 | 24.2 |
| MLHive 3 | No | ND | 34.8 | 21.9 |
| MLHive 4 | No | 35.9 | 34.5 | 23.2 |
| MLHive 5 | No | 35.4 | 31.2 | 23.7 |
| MLHive 6 | No | ND | ND | 23.3 |
| MLHive 7 | No | 30.8 | 27.0 | 23.0 |
| MLHive 8 | No | 36.8 | 33.2 | 23.3 |
| MLHive 9 | No | 36.9 | 33.3 | 22.6 |
| MLHive 10 | Yes | 17.9 | 14.6 | 21.2 |
| MLHive 11 | No | 20.6 | 16.2 | 22.0 |
| MLHive 12 | Yes | 20.2 | 16.3 | 23.4 |
| MLHive 13 | No | 38.2 | 34.3 | 21.2 |
| MLHive 14 | No | 36.7 | ND | 21.7 |
| MLHive 15 | No | 25.8 | 21.5 | 22.3 |
| MLHive 16 | No | ND | 35.3 | 25.0 |
| MLHive 17 | No | ND | ND | 23.7 |
| MLHive 18 | No | ND | ND | 20.9 |
| MLHive 19 | No | 38.0 | 33.7 | 22.0 |
| MLHive 20 | No | ND | ND | 23.4 |
| MLHive 21 | No | ND | 36.2 | 22.9 |
| MLHive 22 | No | ND | ND | 24.2 |
| MLHive 23 | No | 24.1 | 20.1 | 22.3 |

S3: Cq data for hive series ML – *bees* (spore levels in table 2)

**Supplementary figure 4.**
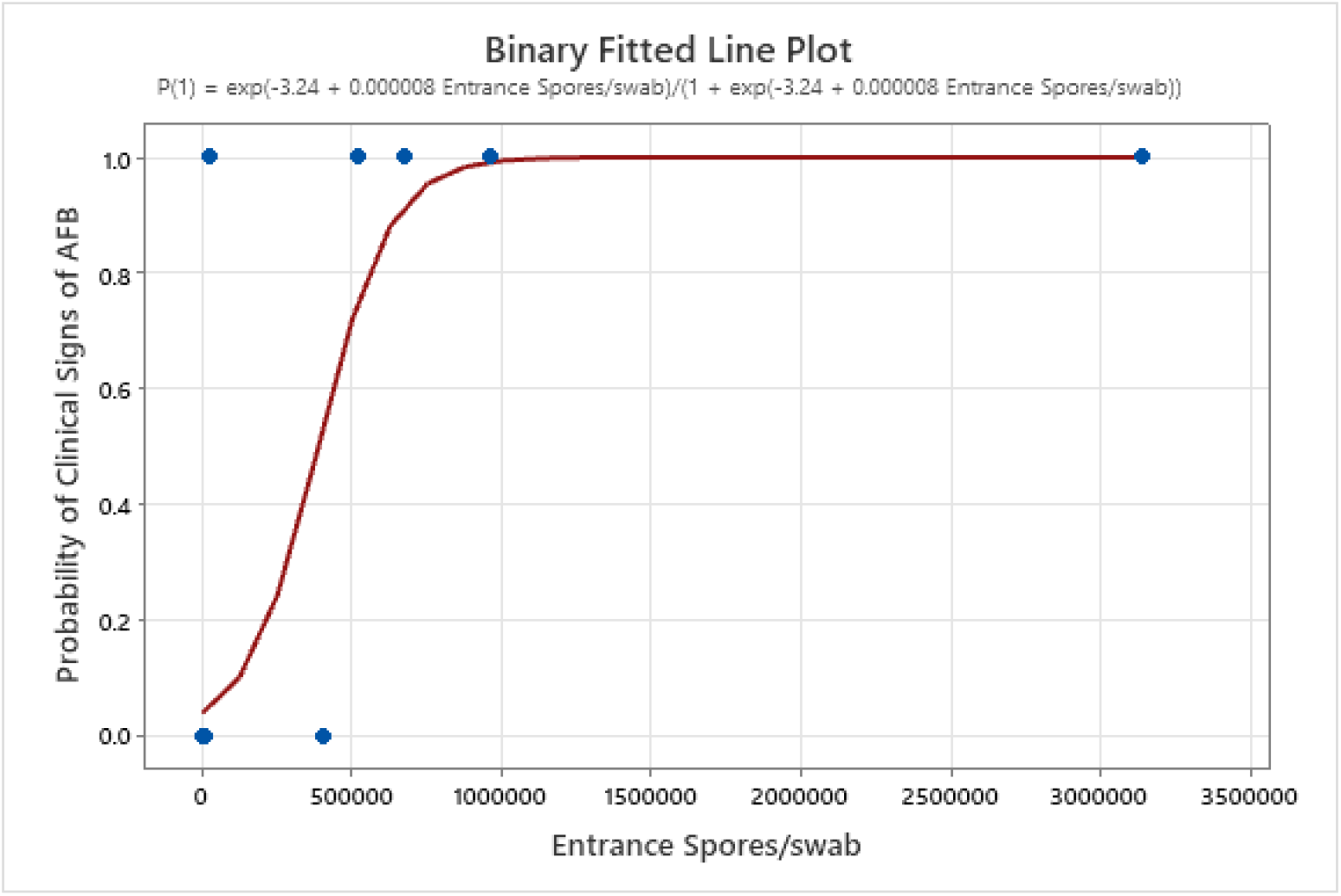
Binary logistic regression model showing the relationship between increasing entrance swab levels and the presence of clinical disease, and the calculated analysis of variance with associated *p*-value (0.000).

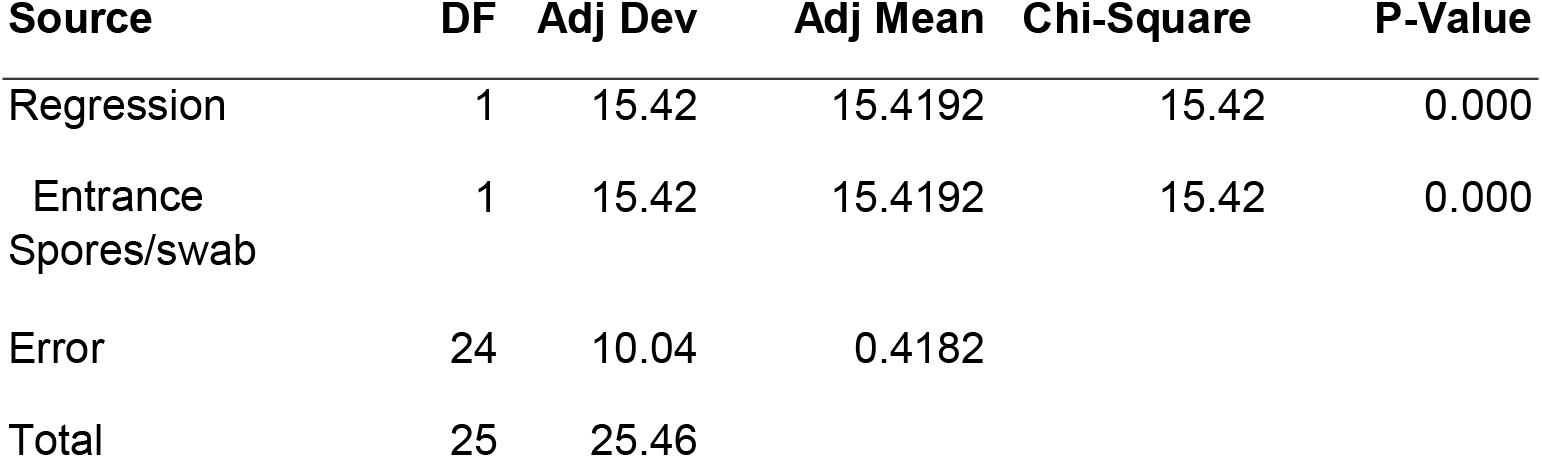
Analysis of variance

| Source | DF | Adj Dev | Adj Mean | Chi-Square | P-Value |
| --- | --- | --- | --- | --- | --- |
| Regression | 1 | 15.42 | 15.4192 | 15.42 | 0.000 |
| Entrance Spores/swab | 1 | 15.42 | 15.4192 | 15.42 | 0.000 |
| Error | 24 | 10.04 | 0.4182 |  |  |
| Total | 25 | 25.46 |  |  |  |

